# Endemic and epidemic human alphavirus infections in Eastern Panama; An Analysis of Population-based Cross-Sectional Surveys

**DOI:** 10.1101/2020.01.10.901462

**Authors:** Jean-Paul Carrera, Zulma M. Cucunubá, Karen Neira, Ben Lambert, Yaneth Pittí, Carmela Jackman, Jesus Liscano, Jorge L. Garzón, Davis Beltran, Luisa Collado-Mariscal, Lisseth Saenz, Néstor Sosa, Luis D. Rodriguez-Guzman, Publio González, Andrés G. Lezcano, Reneé Pereyra-Elías, Anayansi Valderrama, Scott C. Weaver, Amy Y. Vittor, Blas Armién, Juan-Miguel Pascale, Christl A. Donnelly

## Abstract

**Background:** Madariaga virus (MADV), has recently been associated with severe human disease in Panama, where the closely related Venezuelan equine encephalitis virus (VEEV) also circulates. In June, 2017, a fatal MADV infection was confirmed in a community of Darien province.

**Methods:** We conducted a cross-sectional outbreak investigation with human and mosquito collections in July 2017, where sera were tested for alphavirus antibodies and viral RNA. Additionally, by applying a catalytic, force-of-infection statistical model to two serosurveys from Darien province in 2012 and 2017, we investigated whether endemic or epidemic alphavirus transmission occurred historically.

**Results:** In 2017, MADV and VEEV IgM seroprevalence was 1.6% and 4.4%, respectively; IgG antibody prevalences were MADV: 13.2%; VEEV: 16.8%; Una virus (UNAV): 16.0%; and Mayaro virus (MAYV): 1.1%. Active viral circulation was not detected. Evidence of MADV and UNAV infection was found near households — raising questions about its vectors and enzootic transmission cycles. Insomnia was associated with MADV and VEEV infection, depression symptoms were associated with MADV, and dizziness with VEEV and UNAV. Force-of-infection analyses suggest endemic alphavirus transmission historically, with recent increased human exposure to MADV and VEEV in some regions.

**Conclusions:** The lack of additional neurological cases suggest that severe MADV and VEEV infections occur only rarely. Our results indicate that, over the past five decades, alphavirus infections have occurred at low levels in eastern Panama, but that MADV and VEEV infections have recently increased — potentially during the past decade. Endemic infections and outbreaks of MADV and VEEV appear to differ spatially.

**Author summary:** Prior to 2010, it was believed that the Madariaga virus (MADV) was primarily associated with equine disease. However, an outbreak reported in Panama, in an endemic area where Venezuelan equine encephalitis virus (VEEV) also circulates, suggested a change in its epidemiological profile. We aimed to reconstruct the epidemiological dynamics of MADV and VEEV, as well as additional alphaviruses known to circulate in the region in order to understand MADV emergence. For this, cross-sectional serosurveys were used to demonstrate that the Alphaviruses MADV, VEEV and Una virus have repeatedly infected humans in eastern Panama over the past five decades. Whilst their historical transmission has been low, we confirm that the transmission has recently increased for both MADV and VEEV.

## Introduction

Alphaviruses (*Togaviridae: Alphavirus*) are important zoonotic, single-stranded RNA arthropod-borne viruses associated with febrile, severe and sometimes fatal disease in the Americas[1]. Among the most important alphaviruses are eastern equine encephalitis (EEEV), and Venezuelan equine encephalitis viruses (VEEV), and members of the Semliki Forest antigenic complex. These viruses have caused explosive epidemics of human encephalitis and arthritogenic disease in Latin American tropical regions[2,3].

EEEV has recently been reclassified as two different species: EEEV in North America and Madariaga virus (MADV) in other parts of Latin America[4] — each with different predispositions to cause human disease[5]. In 2010, we reported severe neurologic disease in humans associated with MADV infection in Panama[6]. The mechanism underlying this outbreak remains unknown, but age-specific seroprevalence data obtained during the 2010 and 2012 studies suggest recent MADV emergence in Panama[7,8].VEEV is a cause of encephalitis and other pathologies of the central nervous system that can lead to death in humans and domesticated animals in the Americas. This virus causes explosive human and equine epidemics/epizootics, which occur chiefly in South and Central America[9], where the enzootic cycle involves *Culex* mosquitoes (subgenus *Melanoconion)* and sylvatic rodents[10]. Sometimes VEEV causes epizootic outbreaks due to viral adaptations for infection of equids and mosquitoes that allow it to spread rapidly among human and animal populations[11].

The Semliki Forest alphavirus complex includes Mayaro virus (MAYV) and UNAV, which are mostly found in the Amazon region of Peru, Brazil and Venezuela and are characterized by fever and arthralgia, the latter which can persist for years [12]. In the Americas, sizeable human MAYV outbreaks have most often been reported in the Amazon Basin, although, recently this virus was isolated from a febrile child in Haiti, suggesting it may be moving beyond its established territory[13]. UNAV has been detected at low levels during epidemiological studies and surveillance[14,15] but, because this virus has rarely been associated with human disease, the risk to people living in endemic Latin America remains unclear[16]. Both MAYV and UNAV are vectored by forest mosquitoes: *Haemagogus janthinomys* mosquitoes are the primary vectors of MAYV[16], while *Psorophora ferox* and *Psorophora albipes* mosquitoes are thought to be the main vectors of UNAV[17,18]. The MAYV enzootic cycle is also known to involve non-human primates as amplification hosts[16,19].

In June 2017, a fatal MADV infection was confirmed in the Mogue community in Darien, the most eastern province of Panama, prompting field investigations. Here, we use seroprevalence data collected during this survey to determine population exposure and to characterize factors associated with sero-prevalence for MADV and other alphaviruses. By combining seroprevalence survey data from 2012 with that from the recent survey, we also attempted to determine whether alphaviruses emerged recently or were present historically.

## Materials and methods

We reconstructed the epidemiological dynamics of MADV and VEEV using data from cross-sectional surveys undertaken in 2012 and 2017 in Darien Province villages (Figure1). We also identified factors associated with alphavirus exposure, measured as IgG seroprevalence. Map were constructed using the GPS coordinates collecting during the investigation using ArcGIS package online version (Argis Solutions, Inc, Denver, Colorado). Lan use shapes were validate buy the Ministry of Environment (https://www.miambiente.gob.pa)

**Figure 1.**
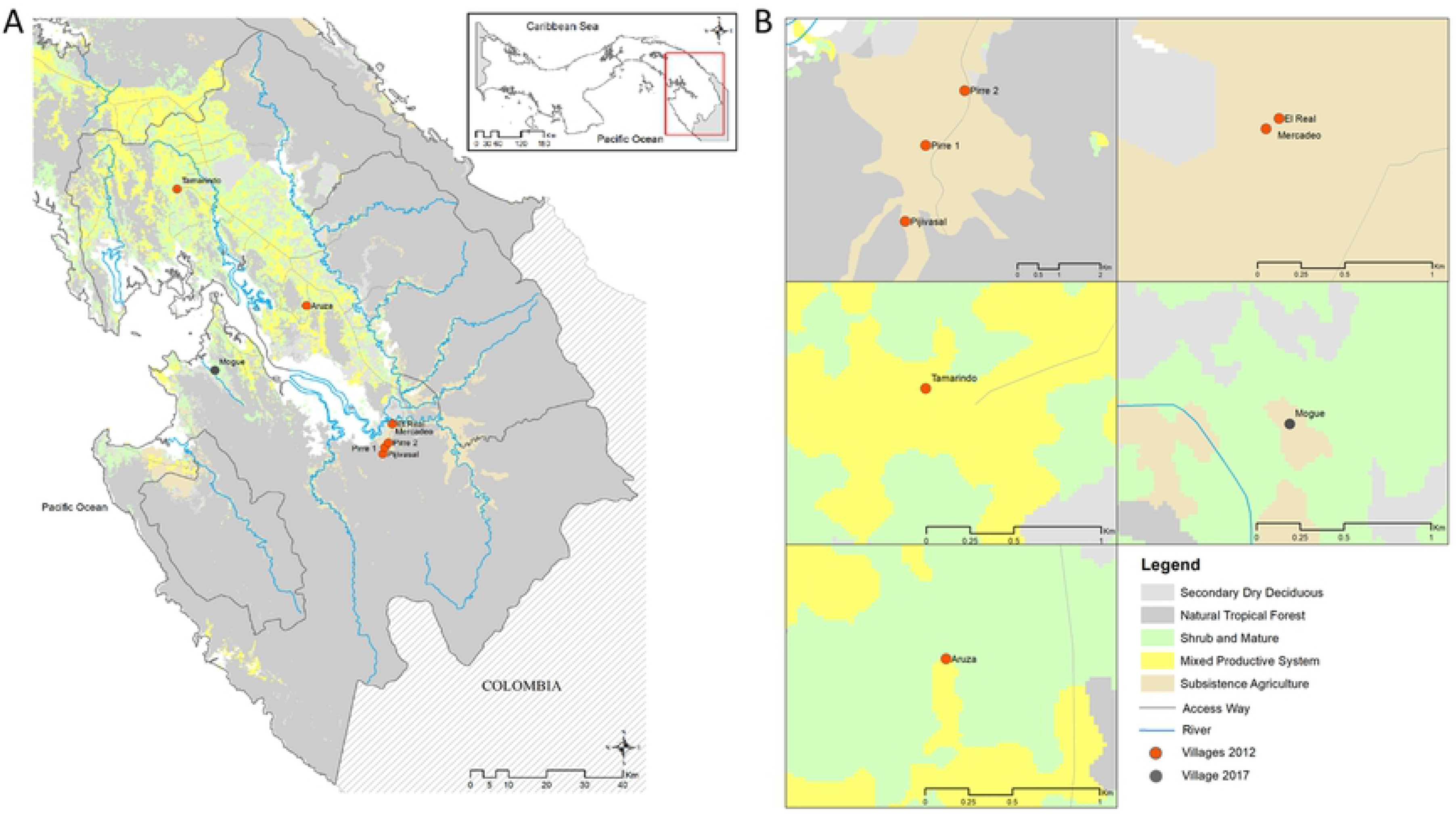
Map of the study sites in eastern Panama: ***A***, the sampling sites in the Darien Province in Eastern Panama. ***B***, Zoom-in projection of sampling sites on a land-use layer

### 2012 Sero-survey

The original 2012 study was conducted by the Gorgas Memorial Institute of Health Studies (GMI) to estimate prevalence and to identify risk factors for zoonotic diseases in Panama[8]. The study included five villages (Figure 1). A total of 897 participants was surveyed but only 774 sera were available for laboratory testing. In Tamarindo, 176 participants were surveyed; 167 in Aruza, 250 in El Real; 130 in Mercadeo; and 174 in Pijibasal/Pirre1-2. All available samples were tested to detect neutralising antibodies against MADV and VEEV using a plaque reduction neutralization test (PRNT). Details of this survey have been described previously[8]. Specific characteristics of the study sites are given in the Supplementary Materials.

### 2017 Sero-survey

On June 30, 2017, a fatal human MADV case was confirmed with viral isolation in Mogue village (Figure 1). This was followed by a collaborative initiative between the Panamanian Ministry of Health and the GMI for outbreak investigation and response. From July 18-22, 2017, 83.3% of inhabitants (250 of 300) were surveyed, including members from all households. Each participant was interviewed using a standardized epidemiological form to record occupation, activities, livestock and crop holdings. Other details are given in the Supplementary Materials and Figure S1. Human sera collected in 2017 were tested using alphavirus genus-specific RT-PCR[20] and by enzyme-linked immunosorbent assays (ELISAs) to detect IgM and IgG antibodies against MADV and VEEV. Positive sera were then confirmed using PRNT with the same method as in the 2012 sero-survey[8]. ELISA antigens were prepared from EEEV-(prepared by Robert Shope at the Yale Arbovirus Research Unit in August 1989) and VEE complex virus (strain 78V-3531)-infected mouse brain. For PRNT, we used chimeric SINV/MADV — shown to be an accurate surrogate for MADV in these assays[21] — and VEEV vaccine strain TC83. In addition, sera were tested for MAYV, UNAV and CHIKV by PRNT using wild type strains (MAYV ARV 0565, UNAV-BT-1495-3 and CHIKV-256899). PRNT80 was positive to more than one virus at a titer of ≥1:20 and there was less than a 4-fold difference in titers.

### Mosquito collection and testing in 2017

Mosquitoes were collected during two consecutive days in Mogue from July 19 to 21 using ten traps: five CDC light traps were baited with octanol, and five Trinidad Traps were baited with laboratory mice. Traps were placed outdoors in peridomestic areas at the edge of the vegetation, from 18:00 to 06:00. Trapped mosquitoes were collected early in the morning and placed in cryovials for storage in liquid nitrogen and transportation to the GMI. Mosquitoes were maintained cold, sorted to species level using taxonomic keys [22] and grouped into pools of 20 individuals.

Mosquito pools were homogenized in 2 mL of minimum essential medium supplemented with penicillin and streptomycin, and 20% fetal bovine serum using a TissueLyser (Qiagen, Hidden, Germany). After centrifugation of 12000 rpm for 10 mins, 200 μL of the supernatant were inoculated in each of two 12.5-cm^2^ flasks of Vero cells. Samples were passaged twice for cytopathic effect (CPE) confirmation. The original mosquito suspensions were used for RNA extraction and tested using alphavirus genus-specific RT-PCR[20].

## Statistical methods

### Associated symptoms and risk factors analysis

We conducted separate analyses for MADV, VEEV and UNAV; in each case, the outcome variable was the presence/absence of antibodies against the virus, as determined by a PRNT_80_ titre ≥ 1:20. The associations between each outcome and self-reported symptoms in the last two weeks were tested using chi-squared and Fisher exact tests; p< 0.05 was considered significant. The associations between each outcome and independent variables were estimated using generalized estimating equations for logistic regression models[23] and were expressed as Odds Ratios (ORs). The most parsimonious model was obtained with log Likelihood Ratio Test (LRT) variable selection[24]. Univariable and multivariable ORs were calculated with 95% confidence intervals.

### Force-of Infection Analysis

To investigate the endemicity and/or recent emergence of three alphaviruses (VEEV, MADV and UNAV), we combined age-structured sero-prevalence data from both surveys (i.e. from 2012[8] and 2017), which encompassed seven sites (Pirre1-2 & Pijibasal, Mercadeo Tamarindo, El Real, Aruza and Mogue) where either human or equine cases of VEEV or MADV have occasionally been reported. See Figure 1 and supplementary materials for a detailed description of these sites.

The historical force-of-infection (FOI) was estimated using a catalytic model[25], where the number of seropositive individuals in each sample was modelled using a binomial distribution,

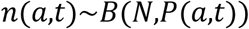

Here *n*(*a*,*t*) is the number of seropositive individuals and *P*(*a*,*t*) is the underlying seroprevalence; in both cases, *a* denotes age and *t* denotes time; *N* is sample size. By making assumptions about *P*(*a*,*t*) (described below), we tested whether MADV, VEEV and UNAV transmission rate has historically been constant over time (“constant FOI’’ model) or has varied— for example, due to recent introduction of these viruses (“time-varying FOI’’ model).

For a constant FOI (λ), we modelled sero-prevalence for age *a* in year *t* (i.e. the time when the sero-survey occurred) as,

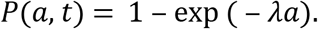

For a time-varying FOI (*λ*_*t*_), we modelled sero-prevalence for age *a* as,

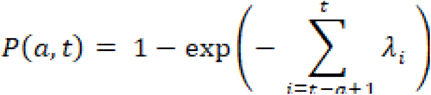

In this framework, we assume no sero-reversion (loss of antibodies over time), no age dependence in susceptibility or exposure [26], and that mortality rate of infected individuals is the same as for susceptible individuals. The models were estimated in a Bayesian framework using Stan’s No-U-Turn Sampler [27,28]. Details of priors and model simulations and packages used are provided in Supplementary Materials. Median of the posteriors distribution of the parameters and their corresponding 95% Credible Intervals (95%CrI) are presented.

### Ethics

The outbreak investigation was undertaken during a public health outbreak response and Ethical approval for use of surveillance data and cross-sectional surveys was given by the GMI Ethics Committee (IRB # 0277/CBI/ICGES/15 and IRB # 047/CNBI/ICGES/11). The written informed consent of participants was obtained. All identifying information of participants was removed and confidentiality was strictly respected. The animal component of this study was approved by the GMI Committee of Care and Use of Animals (001/05 CIUCAL / ICGES, July 4, 2005), and conducted in accordance with Law number 23 of January 15, 1997 (Animal Welfare Guarantee) of the Republic of Panama.

## RESULTS

### Characteristics of the study population

In 2017, 250 participants belonging to 59 houses were surveyed, with complete risk factor data available for only 243 individuals (97.2%). Ages ranged from 1–97 years, and females comprised 51% of surveyed individuals. Further characteristics of the surveyed population are given in Table 1.

**Table 1.** Characteristics of the 243 study participants with complete data from the 2017 survey

| Characteristic | N (%) |
| --- | --- |
| Sex |  |
| Male | 120 (49.4) |
| Female | 123 (50.6) |
| Ages (years) |  |
| 2 – 11 | 80 (32.9) |
| 12 - 30 | 82 (33.7) |
| ≥31 | 81 (33.3) |
| House members* | 4 (2 - 6) |
| **Activities** |  |
| Main occupation |  |
| Student | 122 (50.2) |
| Farmer/Rancher | 48 (19.8) |
| Homemaker/Occupation at home | 73 (30.0) |
| Breeding poultry | 45 (18.5) |
| Fishing for consumption | 7 (2.9) |
| Contact with pastures | 78 (32.1) |
| Contact with crops | 123 (50.6) |
| Clearing vegetation | 80 (32.9) |
| Working in agriculture | 86 (35.4) |
| Working in pastures | 24 (9.9) |
| Working in grain deposits | 21 (8.6) |
| Working in sawmills/forest | 33 (13.6) |
| Working in chicken coops | 58 (23.9) |
| Working in pigsties | 44 (18.1) |
| Washing clothes in ravines or rivers | 111 (45.7) |
| Taking baths in natural water source | 211 (86.8) |
| **House-level features** |  |
| Total houses | 59 |
| House floor material |  |
| Wood | 55 (93.2) |
| Other | 4 (6.8) |
| House with walls | 29 (49.2) |
| House window material |  |
| Concrete (ornamental blocks) | 42 (71.2) |
| Wood | 17 (28.8) |
| Roof material house |  |
| Tin roof | 28 (47.5) |
| Straw thatched | 31 (52.5) |
| Vegetation around the house | 25 (42.4) |
| Rice cultivation around the house | 4 (6.8) |
| Corn cultivation around the house | 3 (5.1) |
| Waste disposal methods |  |
| Burying | 5 (8.5) |
| Burning | 43 (72.9) |
| Other | 11 (18.6) |
| Rain water | 57 (96.6) |

\* range

In 2012, a total of 826 participants was surveyed, but only 774 sera were available for laboratory testing. The risk factors determined from this sero-survey have previously been published [8].

### Alphavirus detection and seroprevalence in 2012 and 2017

In 2012, the overall neutralising antibody seroprevalence was 4.8% (95% CI: 3.4-6.5%) for MADV, and 31.6% (95% CI: 28.3-35.0%) for VEEV.

In 2017, the overall neutralising antibody seroprevalence was: MADV: 13.2% (95% CI 9.2-18.0%); VEEV: 16.8% (95% CI 12.4-22.0%); UNAV: 16.0% (95% CI 11.7-21.1%); and MAYV: 1.2% (95% CI 0.3-3.5%). No evidence of CHIKV infection was found. Neutralising antibody seroprevalence to more than one virus were observed in 3.6% (95% CI 1.6-6.7%) of participants. The proportion of subjects with both MADV and VEEV antibodies was 3.7% (df= 1; Pearson chi2= 3.43;test for independence P= 0.064); both UNAV and VEEV antibodies 3.7% (df=1; Pearson chi2=0.91; test for independence P=0.340) and both MADV and UNAV antibodies 2.9% (df=1; Pearson Chi2= 0.97; test for independence P= 0.325). Only one subject presented antibodies against these three viruses. IgM prevalence was: MADV 1.6% (95% CI 0.4-4.2%); and VEEV 4.4% (95% CI 2.2-7.8%). Concurrent MADV and VEEV IgM was observed in 0.8% of individuals (95% CI 0.1-2.9%). Viral RNA was not detected in sera.

### Associated symptoms and risk factors

Exposure to MADV was significantly associated with self-reported dizziness, fatigue, depression, and difficulty cooking. Having VEEV neutralising antibodies was associated with dizziness and insomnia (Table 2). Participants over 11 years of age were more likely to test positive for UNAV antibodies, with those over 30 years of age being the most likely (Tables 3 and 4). Having a house with walls reduced the risk of testing positive for UNAV antibodies (Tables 3 and 4). The most parsimonious multivariable model revealed that being older and having vegetation around the house were positively associated with MADV antibody prevalence (Table 4). Washing clothes in ravines or rivers was also positively associated with VEEV antibodies in the multivariable model (Table 4).

**Table 2.** Symptoms and signs associated with UNAV, MADV and VEEV exposure (neutralising antibodies)

| Symptoms |  | UNAV+ |  | MADV+ |  | VEEV+ |  |
| --- | --- | --- | --- | --- | --- | --- | --- |
|  | N (%) | n(%)** | ***P value | n(%)** | ***P value | n(%) | ***P value |
| Fatigue | 85 (35.0) | 14 (35.0) | 0.998 | 15 (48.4) | 0.094 | 19 (45.4) | 0.125 |
| Difficulty with concentration | 60 (24.7) | 13 (32.5) | 0.210 | 10 (32.3) | 0.296 | 11 (26.2) | 0.804 |
| Memory loss | 58 (23.9) | 12 (30.0) | 0.320 | 11 (35.5) | 0.104 | 13 (31.0) | 0.236 |
| Confusion | 41 (16.9) | 10 (25.0) | 0.133 | 6 (19.4) | 0.693 | 11 (26.2) | 0.076 |
| Dizziness | 72 (29.6) | **18 (45.0)** | **0.020** | 12 (38.7) | 0.236 | **18 (42.9)** | **0.039** |
| Seizures | 5 (2.1) | 2 (5.0) | 0.191 | 2 (6.5) | 0.123 | 2 (4.8) | 0.207 |
| General weakness | 65 (26.7) | 15 (37.5) | 0.093 | **13 (41.9)** | **0.041** | 13 (31.0) | 0.499 |
| Paralysis | 11 (4.5) | 3 (7.5) | 0.396 | 1 (3.2) | 1.000 | 4 (36.4) | 0.102 |
| Difficulty ambulating | 29 (11.9) | 5 (12.5) | 0.540 | 5 (16.1) | 0.302 | 8 (19.1) | 0.118 |
| Headache | 110 (45.3) | 22 (55.0) | 0.176 | 15 (48.4) | 0.709 | 21 (50.0) | 0.498 |
| Insomnia | 33 (13.6) | 3 (7.5) | 0.313 | **9 (29.0)** | **0.012** | **12 (28.6)** | **0.002** |
| Depression | 22 (9.1) | 5 (12.5) | 0.285 | **6 (19.4)** | **0.044** | 2 (4.8) | 0.228 |
| Irritability | 16 (6.6) | 3 (7.5) | 0.732 | 2 (6.5) | 1.000 | 4(9.5) | 0.490 |
| Difficulty cooking | 23 (9.5) | 5 (12.5) | 0.473 | **6 (19.4)** | **0.044** | 6 (14.3) | 0.241 |
| Difficulty cleaning | 28 (11.5) | 5 (12.5) | 0.832 | 6 (19.4) | 0.144 | 5 (11.9) | 0.932 |
| Difficulty working | 25 (10.3) | 3 (7.5) | 0.776 | 6 (19.4) | 0.075 | 6 (14.3) | 0.348 |
| Fever | 6 (2.5) | 1 (2.5) | 1.000 | 0 (0.0) | 1.000 | 1 (2.4) | 0.173 |
| Chills | 2 (0.8) | 1 (2.5) | 0.303 | 0 (0.0) | 1.000 | 0 (0.0) | 1.000 |
| Emesis | 1 (0.4) | 0 (0.0) | 1.000 | 0 (0.0) | 1.000 | 1 (2.4) | 0.173 |
| Diarrhea | 1 (0.4) | 0 (0.0) | 1.000 | 0 (0.0) | 1.000 | 1 (2.4) | 0.173 |

n=40 with UNAV antibodies; n=31 with MADV antibodies; n=42 with VEEV antibodies n=243 participants in total\* Overall proportion of participants with symptoms \*\* Proportion of those with antibodies that reported symptoms \*\*\*results with p < 0.05 are shown in boldface type +based on PRNT results

**Table 3.**
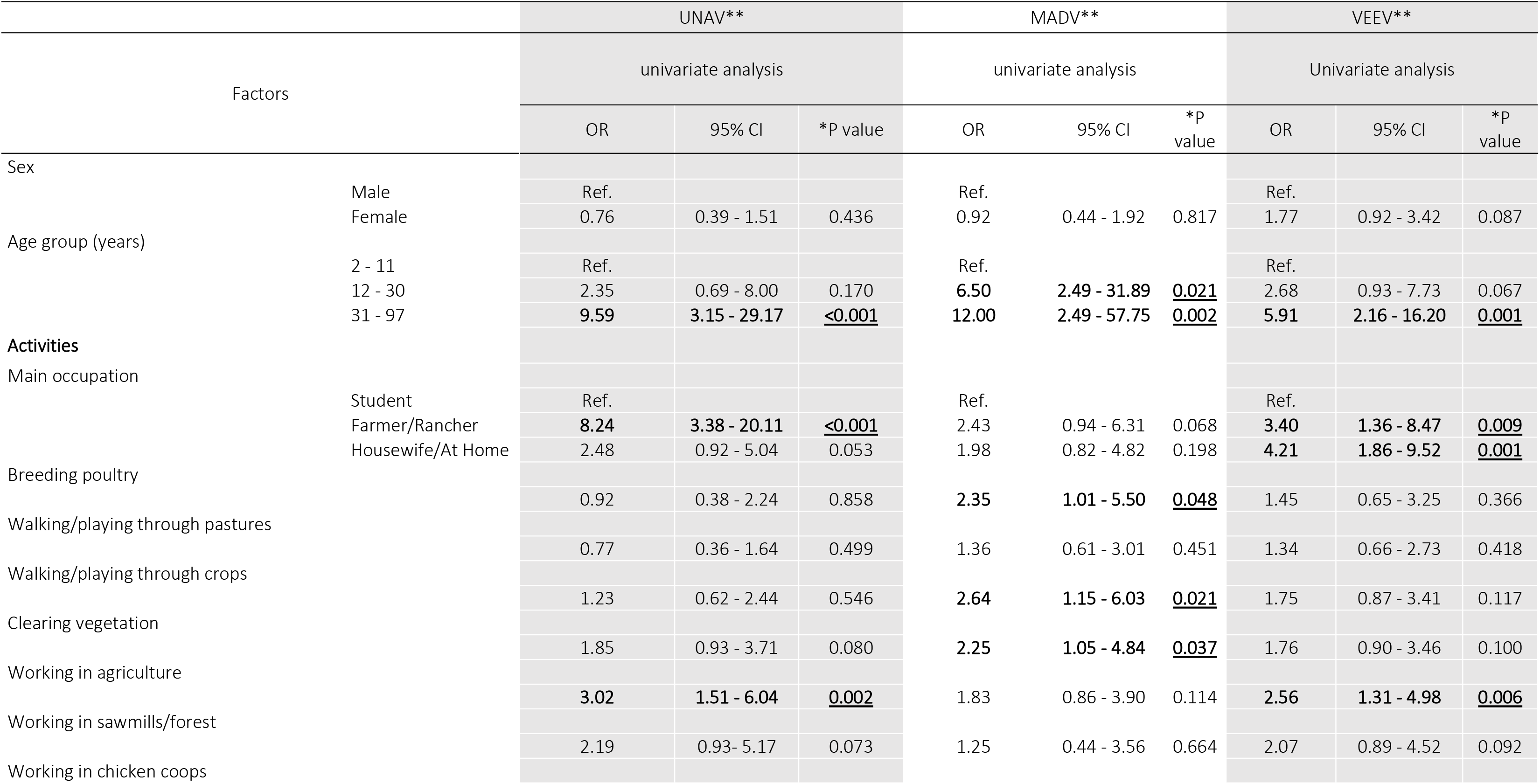
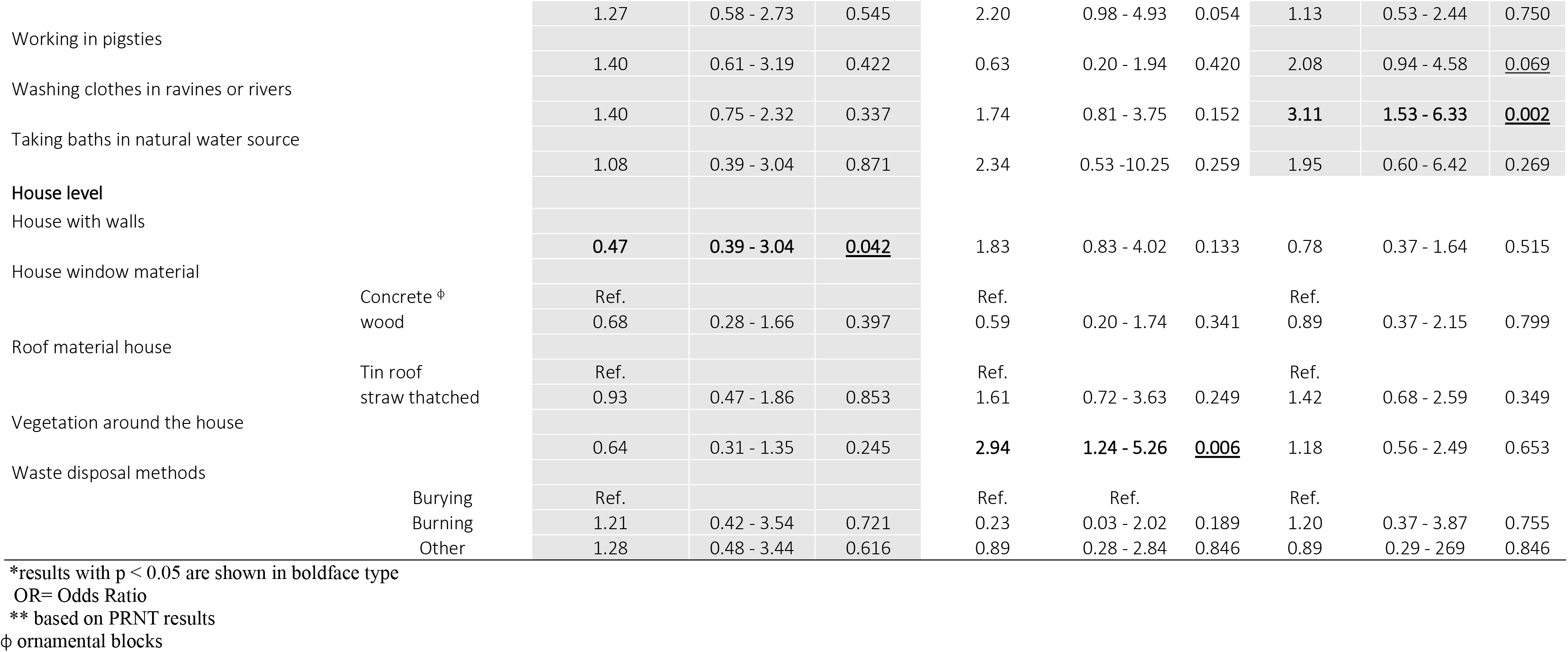
Independent factors associated with the seroprevalence of UNAV, MADV and VEEV neutralising antibodies in univariate generalized estimating equations for logistic regression models (n=243)

**Table 4.** Independent factors associated with the seroprevalence of UNAV, MADV and VEEV neutralising antibodies in multivariable generalized estimating equations for logistic regression models (n=243)

|  |  | UNAV* |  |  | MADV* |  |  | VEEV* |  |  |
| --- | --- | --- | --- | --- | --- | --- | --- | --- | --- | --- |
|  |  | Multiple regression |  |  | Multiple regression |  |  | Multiple regression |  |  |
| Factors |  | OR | 95% CI | **P value | OR | 95% CI | **P value | OR | 95% CI | **P value |
| Age group (years) |  |  |  |  |  |  |  |  |  |  |
|  | 2 - 11 | Ref. |  |  | Ref. |  |  | Ref. |  |  |
|  | 12 - 30 | 2.39 | 0.70 - 8.15 | 0.164 | **6.28** | **1.27 - 31.00** | **<u>0.024</u>** | 1.83 | 0.61 - 5.53 | 0.279 |
|  | ≥31 | **9.98** | **3.27 - 30.48** | **<u><0.001</u>** | 12.64 | 2.61 - 60.19 | **<u>0.002</u>** | 4.53 | **1.61 - 12.74** | **<u>0.004</u>** |
| House with walls |  |  |  |  |  |  |  |  |  |  |
|  |  | **0.43** | **0.20 - 0.94** | **<u>0.035</u>** |  |  |  |  |  |  |
| Washing clothes in ravines or rivers |  |  |  |  |  |  |  |  |  |  |
|  |  |  |  |  |  |  |  | 2.65 | 1.24 - 5.63 | **<u>0.011</u>** |
| Vegetation around the house |  |  |  |  |  |  |  |  |  |  |
|  |  |  |  |  | **2.96** | **1.25 - 6.98** | **<u>0.013</u>** |  |  |  |

\* based on PRNT results \*\*results with P < 0.05 are shown in boldface type OR= Odds Ratio

### Enzootic vectors

In 2017, a total of 113 mosquitoes across ten species was collected: *Culex* (*Culex*) *coronator* (36.3%), *Cx.* (*Melanoconion*) *pedroi* (14.2%), *Cx.* (*Mel.*) *spissipes* (10.6%), *Cx.* (*Cx.*) *nigripalpus* (10.6%), *Cx.* (*Mel.*) *vomerifer* (8.8%), *Cx.* (*Cx.*) *declarator* (5.3%), *Cx.* (*Mel.*) *adamesi* (2.7%), *Cx.* (*Mel.*) *dunni* (2.7%). The overall mean number of females per trap-night was 6.7 in the Trinidad traps compared with 4.6 in the CDC traps. No viruses were detected in samples from mosquitoes.

### Alphavirus Force-of-Infection

For each virus, we fit both constant and time-varying FOI models to the seroprevalence data (see Methods) to describe the per capita rate at which susceptible individuals become infected per year. Since the constant FOI model is effectively nested within the time-varying FOI model, we report on whether the latter model improved the fit relative to the former.

Our results indicate temporal and geographic heterogeneity in the human population’s exposure to MADV (Figure 2), VEEV (Figure 3) and UNAV (Figure 4). The highest estimated sero-prevalence of each of the three viruses in under-10-year-olds (an indirect metric of recent transmission) were estimated for VEEV in Pirre 1-2 & Pijibasal at a posterior median of 44.8% (95%CrI: 34.9-55.0%), followed by UNAV in Mogue at 5.6% (95%CrI: 4.1 – 7.5) and by MADV in Aruza at 4.7% (95%CrI: 3.2-6.7%).

**Figure 2.**
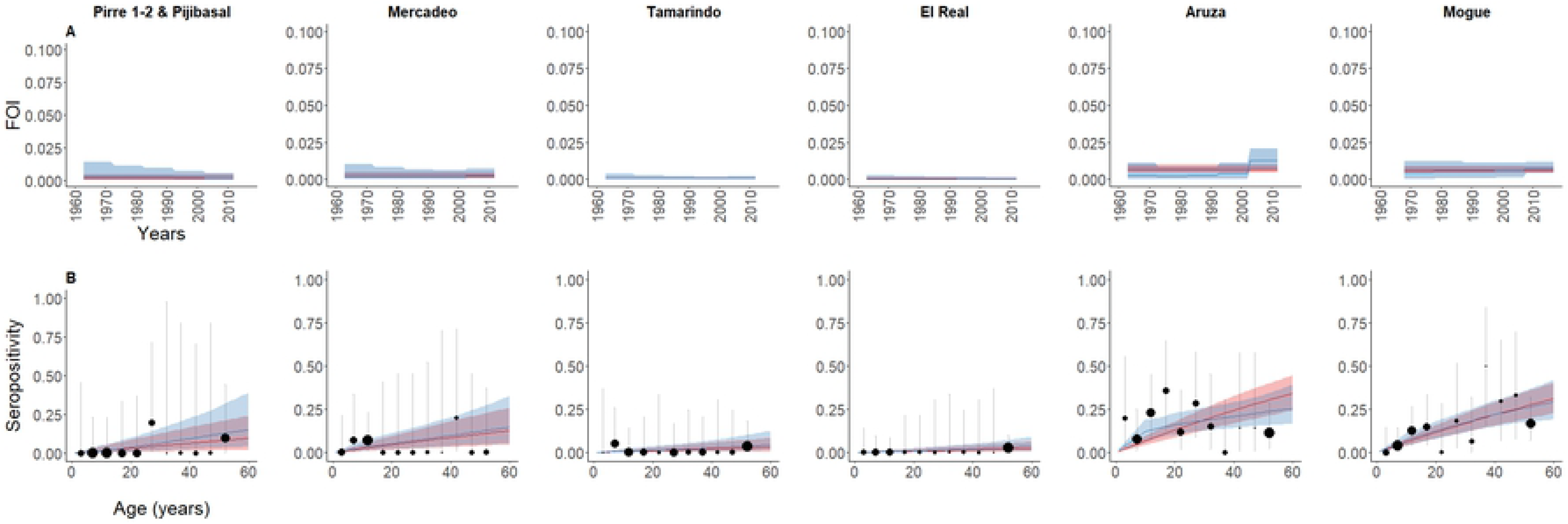
Force-of-Infection (FOI) models fitted to MADV seroprevalence data. ***A*** (top panels), estimated constant (red) vs time-varying force-of-infection (blue) for MADV in eastern Panama over 50 years and ***B*** (bottom panels) fitted and observed seroprevalence. Red lines represent the estimated constant force-of-infection and blue lines the estimated time-varying force-of-Infection. In each case the shading represents 95% credible intervals from the model. The circles’ radii in the lower panels indicates sample size in each 5-year age group and the vertical lines represent 95% confidence intervals for observed seroprevalence.

**Figure 3.**
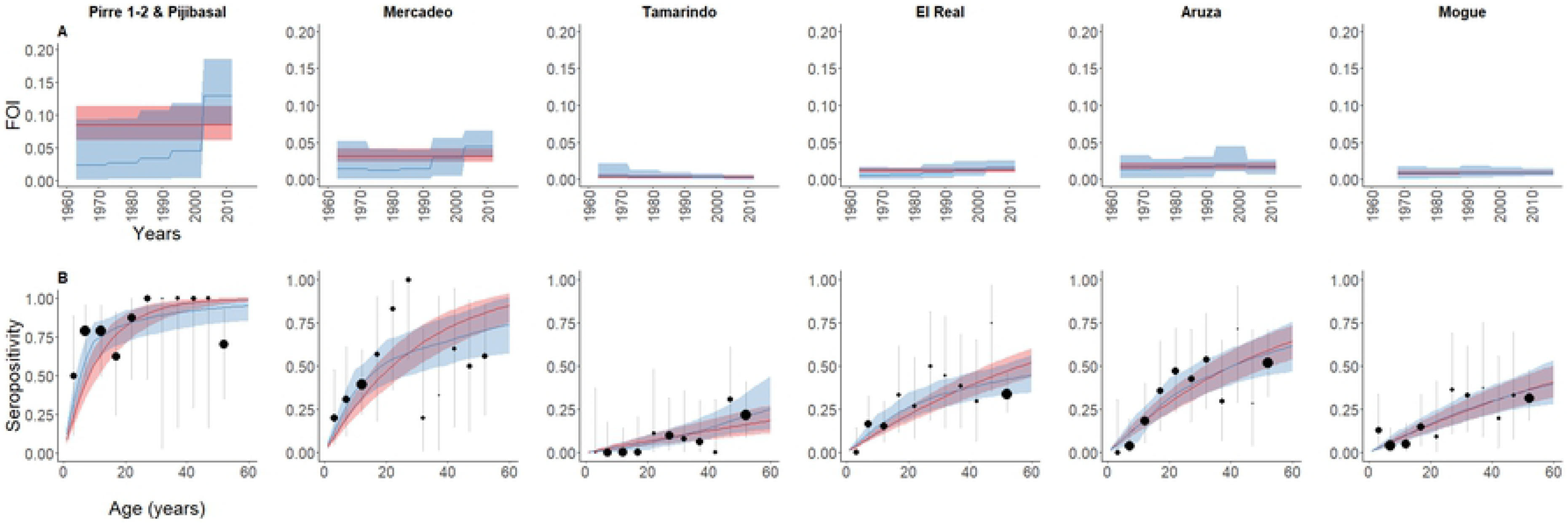
Force-of-Infection (FOI) models fitted to VEEV seroprevalence data. ***A*** (top panels), estimated constant (red) vs time-varying force-of-infection (blue) for VEEV in eastern Panama over 50 years and ***B*** (bottom panels) fitted and observed seroprevalence. Red lines represent the estimated constant force-of-infection and blue lines the estimated time-varying force-of-Infection. In each case the shading represents 95% credible intervals from the model. The circles’ radii in the lower panels indicates sample size in each 5-year age group and the vertical lines represent 95% confidence intervals for observed seroprevalence.

**Figure 4.**
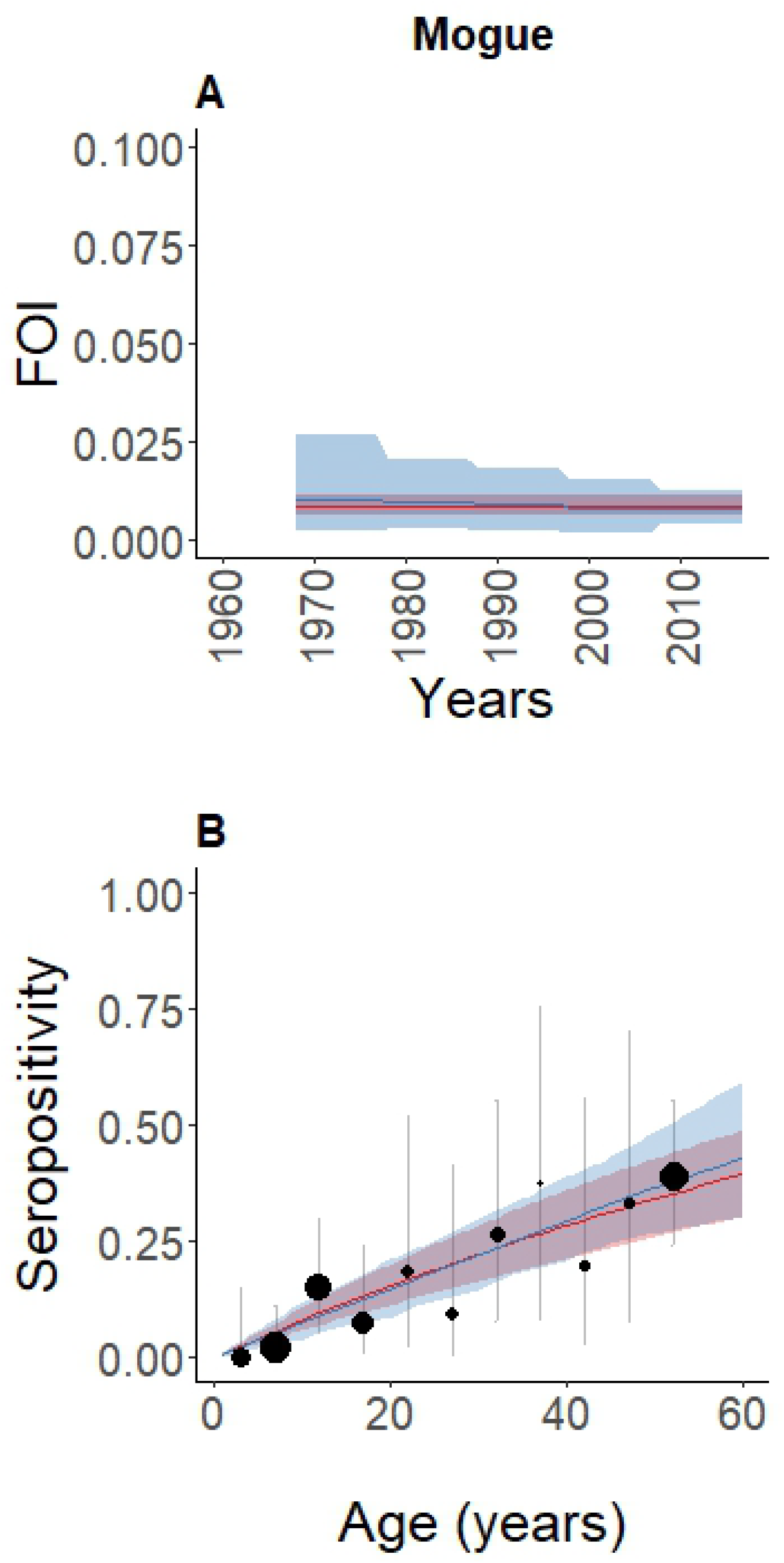
Force-of-Infection (FOI) models fitted to UNAV seroprevalence data. ***A*** (top panels), estimated constant (red) vs time-varying force-of-infection (blue) for UNAV in eastern Panama over 50 years and ***B*** (bottom panels) fitted and observed seroprevalence. Red lines represent the estimated constant force-of-infection and blue lines the estimated time-varying force-of-Infection. In each case the shading represents 95% credible intervals from the model. The circles’ radii in the lower panels indicates sample size in each 5-year age group and the vertical lines represent 95% confidence intervals for observed seroprevalence.

For MADV, in six of the seven locations, there was no evidence of time-varying transmission (Table 5); but in one location, Aruza, FOI was estimated as 0.012 (CrI95% 0.006 – 0.021) (Figure 2A) in the latest decade analyzed (2002-2012) —a multiple of 4.6 and 5.3 times (ratio of posterior medians) the values estimated for 1992-2012 and 1982-1992, respectively (Figure 2B).

**Table 5.**
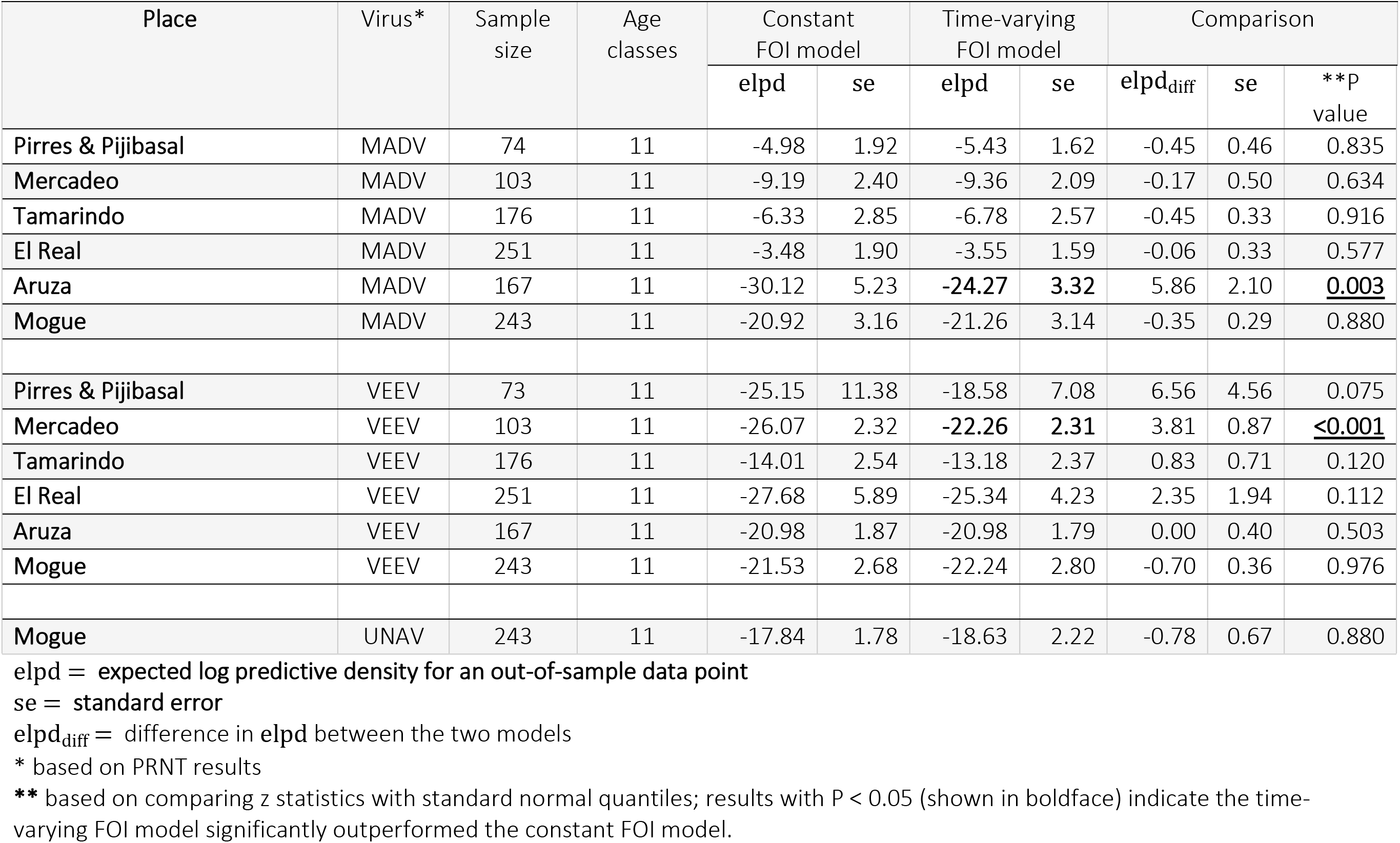
Comparison of constant versus time-varying Force-of-Infection for UNAV, MADV and VEEV in 2012 and 2017

For VEEV, in six of the seven locations, there was no statistical support for time-varying transmission (Table 5). For the constant model, we estimated an annual FOI of 0.08 (95%CrI, 0.06 – 0.11) for VEEV in Pirre 1-2 & Pijibasal corresponding, to seroprevalence reaching 75% in 15-year-olds and almost 100% by 60 years of age (Figure 3A). However, from the relatively small sample (only 75 subjects), it is unclear whether these results are due to consistently high endemic transmission or recent introductions and/or recent outbreaks. For one location, Mercadeo, a time-varying FOI model fit the data best. In this case, FOI in the most recently analyzed decade (2002-2012) was estimated at 0.04 (95%CrI: 0.03 – 0.06) — an increase of 1.5 times (ratio of posterior medians) over the previous decade (1992-2012) and 3.1 times that of 1972-1992 (Figure 3B).

For UNAV, only tested in Mogue, a constant model fit the data best with a FOI estimated at 0.008 (95% CrI: 0.006-0.011) (Figure 4).

## Discussion

By analyzing data from recent cross-sectional seroprevalence studies, we reconstructed alphavirus transmission in eastern Panama. Historical transmission rates indicated endemic transmission of VEEV, MADV and UNAV in humans with increased human exposure during the past decade. Here, we show evidence of acute IgM antibody responses against MADV and VEEV in people without signs of neurologic disease, suggesting asymptomatic infections or mild disease. To our knowledge, this is the first evidence of human infection with UNAV in Panama, even though its circulation was reported during the 1960s in mosquitoes (*Psorophora ferox* and *Ps. albipes*) collected in western Panama[17]. Our results also demonstrate the highest seroprevalence of UNAV reported in Latin America[15,29].

Using catalytic FOI models fit to age-stratified seroprevalence data, we reconstructed 50 years of historical transmission rates for VEEV and MADV for seven locations in Darien Province. In most locations, the data indicated consistent endemic transmission of these viruses. In two locations – Mercadeo (for VEEV) and Aruza (for MADV) — there was evidence of a recent increase in human exposure. These results suggest that MADV and VEEV incidence differ geographically. The observed FOI profile suggest that VEEV infections increased in Pirre 1-2 & Pijibasal and Mercadeo, locations surrounded by tropical forest, while MADV infections increased mostly in Aruza, a formerly forested area converted to agricultural land over 30 years ago [30]. Although ecological changes could be associated with the increased exposure to MADV in Aruza, it is unclear which drivers could also explain the simultaneous rise in VEEV we estimated.

Only 3.6% of participants had antibodies to more than one alphavirus. Mixed alphavirus antibody responses in Peru[5] and Panama[8] suggest cross-protective immunity. However, the mechanism of cross-protection and whether some alphaviruses induce a stronger heterologous response than others remain unclear.

The MADV seroprevalence in 2017 was greater for those living with vegetation around the house, contrasting with previous evidence in 2012, suggesting possible change in exposure risk[8]. However, characteristics of houses in Mogue in 2017 may differ from areas that were surveyed in 2012[8]. Potential MADV vectors within the *Culex* (*Melanoconion*) subgenus[31] were found during our peridomestic investigation in Mogue. This finding of vectors near houses with surrounding vegetation as a risk factor supports the hypothesis that MADV infections can occur near houses. This contrasts with VEEV risk factors, which include washing clothes in ravines or rivers, suggesting that VEEV seropositivity is associated with human incursion into the gallery forest, a potential natural habitat for development of larvae of the main vectors *Culex* (*Melanoconion*) spp[31].

Having a house with walls was associated with lower UNAV sero-prevalence in Mogue. This suggests that UNAV infections can also occur outside of the forest, where the main vector *Ps. ferox* and non-human primates are believed to maintain the enzootic cycle[16,17,19]. *Psorophora* spp. have been also found in disturbed areas of Panama[32], indicating potential changes in the vector habitat usage.

Alphaviral exposure was associated with several self-reported neurological and constitutional sequelae. Specifically, weakness, insomnia, depression and dizziness were commonly associated with prior MADV, VEEV, and UNAV exposure. Depression and other neurological symptoms have also been observed after neurotropic flavivirus infections in North America[33]. However, the role of several alphaviruses in long-term neurological impairment is still unknown. This highlights the need to further investigate the long term ramifications of alphaviral infection with objective testing (e.g. neuropsychological testing, imaging).

Alphaviral RNA was not detected in samples from either humans or mosquitoes, even though field surveys and collection were performed soon after the confirmation of a fatal MADV infection in the community. Although sample size is always a limiting factor in attempts to identify ongoing infections, these results suggest that these alphaviruses may be short-lived peripherally, or produce low viremia[7]. Low MAYV sero-prevalence was also detected in our earlier research[7], indicating little human exposure to this virus in Panama.

Our study has several limitations. Clinical outcomes statistically associated to exposure to these alphaviruses represent exploratory, and causal inference studies that should be followed up with more comprehensive assessments. Our study only obtained preliminary data during an outbreak response to generate hypotheses. Although mosquito collections were only performed over two days, and the number of collected mosquitos does not allow us to draw conclusions about active viral circulation. The collection of few mosquitos vectors near houses suggest close contact between vectors and humans. The use of both CDC traps baited with octanol and Trinidad traps enhanced our ability to captured alphavirus enzootic vectors [34]. The sample size used in these sero-surveys only allowed us to describe general trends in the force-of-infection over time. Also, we cannot exclude cross-reactivity or age-dependency in exposure or susceptibility. More precise estimates would require an increased sample size and, ideally, longitudinal data collection.

In summary, we investigated alphavirus transmission in Panama using age-specific seroprevalence data to look back over five decades. Our results suggest that human alphavirus infections may have gone undetected by the Panamanian surveillance system, and hint that the MADV and VEEV outbreaks in 2010 may have been due to a common increase in enzootic circulation. The antibody seroprevalence we determined for UNAV is the highest reported in Latin America. Taken together, these results coupled with potential symptoms of MADV and VEEV infection underscore the importance of developing comprehensive arboviral surveillance in Latin American enzootic regions.

## Supporting Information Legends

1. **Supporting information file 1**. S1. Includes detailed information on the study sites, equations, figures and references of the manuscript.
2. **Supporting information file 2**. S2 Checklist: STROBE Checklist

## Acknowledgments

We thank the people from the Mogue community for cooperation and hospitality during our investigation as well as Patricia Aguilar for technical suggestions and support with reagents. We also thank Mileyka Santos for mosquito identification; Isela Guerrero, Jose Francisco Galue, Marisin Tenernorio and Daniel Castillo for technical support with the RT-PCR and ELISAs testing; Sandra Lopez-Verges, for provided reagents and revision of the manuscript. JMP, BA and AV are members of the Sistema Nacional de Investigación (SNI), Panama.

## Financial support

JPC is funded by the Clarendon Scholarship from University of Oxford and Lincoln-Kingsgate Scholarship from Lincoln College, University of Oxford [grant number SFF1920_CB2_MPLS_1293647]. This work was supported by SENACYT [grant number FID-16-201] grant to JPC and AV. Also, the Neglected Diseases Grant from the Ministry of Economy and Finance of Panama to JMP [grant number 1.11.1.3.703.01.55.120]. BA received support from the Panamanian Ministry of Economy and Finance and the Panamanian Ministry of Health [grant number 06-2012-FPI-MEF/056-2012-MINSA]. SCW is supported by the U.S. National Institutes of Health [grant number R24AI120942]. ZMC and CAD acknowledge joint Centre funding from the UK Medical Research Council and Department for International Development [grant number MR/R015600/1]. ZMC is funded by the MRC Rutherford Fund Fellowship [grant number MR/R024855/1]. CAD acknowledge funding some the National Institute of Health Research for support of the Health Protection Research Unit in Modelling Methodology.

## Disclaimers

The opinions expressed by authors contributing to this journal do not necessarily reflect the opinions of the Gorgas Memorial Institute of Health Studies, The Panamanian Government, or the institutions with which the authors are affiliated.

## Potential conflicts of interest

All Authors: No reported conflicts of interest. Conflicts that the editor consider relevant to the content have been disclosed.

